# Social influence in adolescence: behavioral and neural responses to peer and expert opinion

**DOI:** 10.1101/2023.07.06.547708

**Authors:** Fatemeh Irani, Joona Muotka, Pessi Lyyra, Tiina Parviainen, Simo Monto

**Affiliations:** Centre for Interdisciplinary Brain Research, Department of Psychology, University of Jyväskylä, Finland;, University of Jyväskylä, Finland

**Keywords:** Adolescence, normative influence, informational influence, MEG, FRN, beta oscillation

## Abstract

Social influence plays a crucial role during the teen years, with adolescents supposedly exhibiting heightened sensitivity to their peers. In this study, we examine how social influence from different sources, particularly those with varying normative and informational significance, affect adolescents’ opinion change. Furthermore, we investigate underlying neural dynamics to determine whether these two behaviorally similar influences share their neural mechanisms. Twenty-three participants (14-17 years old) gave their opinions about facial stimuli and received feedback from either a peer group or an expert group, while brain responses were recorded using concurrent magnetoencephalography. In a second rating session, we found that participants’ opinions changed in line with conflicting feedback, but only when the feedback was lower than their initial evaluation. On the neural level, conflict with the peer group evoked stronger neural responses than conflict with experts in the 230-400 ms time window. Nevertheless, there was no greater conformity toward peers. Moreover, conflict compared to no conflict decreased neural oscillations in the beta frequency range (20–27 Hz) at the right frontal and parietal channels. Taken together, our findings do not support the general assumption that adolescent behavior is excessively vulnerable to peer norms, although we found heightened neural sensitivity to peer feedback.

## Introduction

Adolescence is characterized by notable transformations in physical, cognitive, emotional, and social aspects (Guyer et al., 2016). During this period, individuals start to navigate broader social environments, as they form their identities and social roles (Blakemore & Mills, 2014). In this stage, adolescents become increasingly sensitive to their peers - those of the same age, status, or skill level (Brown & Larson, 2009). They become especially motivated to maintain alignment with their peer group’s norms and expectations (Berns et al., 2010; Wasylyshyn et al., 2018; Molleman et al., 2022), which is known as normative influence and serves to preserve positive social relations (Deutsch & Gerard, 1955). Normative influence often creates compliance, where individuals publicly conform to the norm while privately maintaining their opinions (Cialdini & Goldstein, 2004). However, it can result in authentic influence and private acceptance when an individual perceives belonging to the group as rewarding or trusts the group members’ judgments (Spears, 2021). Another type of influence affecting individuals’ behavior is detached from identity concerns and approval seeking and is known as informational influence. This also relies on social cues but serves to maximize the accuracy of judgments (Deutsch & Gerard, 1955).

Real-world social settings for adolescents involve both peer interactions and guidance from adults such as teachers, parents, and experts. The impact of peers is often posited to surpass other sources during adolescence (Crone & Dahl, 2012). This tendency toward peer opinions concerns practitioners, public health experts, and parents, because it may place adolescents in a position vulnerable to increased health risk and maladaptive decision-making (Simons-Morton & Farhat, 2010). Crucially, previous neuroimaging studies reporting the significant influence of peers on adolescent behavior have not contrasted the peer effect with other social sources of information. This narrow focus may have limited our understanding of whether the observed effect is specific to peer interactions or due to social influence in general (e.g., Berns et al., 2010; Wasylyshyn et al., 2018). Engelmann and colleagues (2012) conducted a study on the effect of expert advice on decision making in adults and adolescents. The results revealed that in comparison to adults, expert advice had a greater influence on adolescents. These findings imply that adolescents may exhibit heightened sensitivity to all social inputs. This highlights the importance of comparing different sources of influence in adolescence.

Some studies have used common interactions to compare the impact of peers and parents on adolescent’s risk-taking behavior (Van Hoorn et al., 2018), subjective evaluation of artwork (Welborn et al., 2016), and everyday constructive and unconstructive behavior (Do et al., 2020). These studies aimed to disentangle the effects specific to each source. Unexpectedly, behavioral findings revealed no distinct effect of peers and parents on behavior. Neurally, the BOLD signal from fMRI measurement revealed greater activity in reward-related, mentalizing, and cognitive control regions in the presence of peers (Van Hoorn et al., 2018; Do et al., 2020) and greater functional connectivity in these regions in the presence of parents (Van Hoorn et al., 2018). These findings challenged the assumption of adolescents orienting away from the family and toward peers and may be attributed to the perception of parents as an authoritative and normative source rather than robust informational sources.

In an investment game study with adult participants, advice was provided by a financial expert and a peer. Despite the advice being randomized to be identical for the expert and peer, individuals followed the expert’s advice longer. The anterior cingulate cortex (ACC) and the superior frontal gyrus showed more activation when participants chose to ignore the expert’s advice compared to ignoring the peer’s advice (Suen et al., 2014). The activation of the ACC regions is linked to performance monitoring and error detection. It is typically observed during a disagreement with the group in social influence contexts and is presumed to guide future action and behavior modification (Klucharev et al., 2009; Berns et al., 2010). In Suen and colleagues’ study, the activation of the ACC during ignoring the advice of the expert is considered an indicator of error detection and signals the need to adjust future behavior based on the expert’s advice. With the adolescent population, following the peer for a longer time may occur when both peer and expert opinions are presented. However, in a behavioral probabilistic learning task, with advice provided by both peers and older adults, adolescents showed a preference for stimuli recommended by the older adult rather than conforming to their peer group, contrary to the assumption of peer sensitivity. This suggests that in certain decision-making contexts, adolescents prioritize the advice of older adults over that of their peers (Lourenco et al., 2015).

While many studies have demonstrated peer influence on adolescent behavior, studies with direct comparison have indicated that peers do not always outweigh other sources of influence. The current study aims to examine whether social influence from different sources has different effects on adolescents’ behavioral and neural responses. We manipulated social influence in a rating task using two sources with different normative and informational weights: a peer group and an expert group. We hypothesized that adolescents conform more to peers than experts when receiving conflicting feedback.

To identify the neural processes underlying conformity behavior to different sources, we used magnetoencephalography (MEG) to capture neural markers of conflict between individual and group opinions. MEG/EEG studies on social influence have shown that a frontocentral evoked component at ∼200 ms latency known as feedback-related negativity (FRN) reflects the response to group conflict, similar to the activation of ACC and deactivation of ventral striatum, which are associated with conflict monitoring and reward processing, respectively. Thus, we hypothesized that a discrepancy between an individual’s preference and the peer group’s opinion would evoke stronger FRN-like neural responses compared to a discrepancy with the expert opinion (Kim et al., 2012; Shestakova et al., 2013; Zubarev et al., 2017). In addition to evoked responses, it has been observed that when there is a disagreement between an individual’s opinion and the group’s opinion, the associated cognitive and emotional responses are manifested in modulations of neuronal oscillatory activity, specifically theta (4-8 Hz) synchronization and beta (13-30 Hz) desynchronization (Zubarev et al., 2017; Irani et al., 2022). Consequently, we predicted that conflict with peer group opinion would be followed by an increase in the power of theta band oscillations and a decrease in the power of beta-band oscillations compared to conflict with experts.

## Methods

### Participants

Twenty-three female participants (mean age: 16.08 years, SD: 1.08, 21 right-handed) with no reported history of psychiatric or neurological illnesses took part in the study. All participants had a normal or corrected-to-normal vision. The study was approved by the ethical committee of the University of Jyväskylä, and all participants and their guardians signed informed consent. Participants were compensated with a 15-euro grocery gift card.

### Stimuli and task

We employed an adapted version of the evaluative judgment task (Klucharev et al., 2009). Visual stimuli with the size of 19×25 cm was shown in the middle of the screen, positioned 1 meter from the participants, and presented with a DLP projector. The stimuli were 240 female faces in the age range of 18-35, from free internet sources which were modified to look younger using the FaceApp photo editing application (FaceApp Technology Limited). A separate group of 10 females (mean age: 21.60, SD: 3.89) rated the age of a subset of the modified photos (n=40) and the estimated ages were within the intended age range of 14-17. None of the individuals reported the photos as manipulated.

For the MEG experiment, the participants were instructed to rate the sociability of the faces on an 8-point scale (1=least sociable, 8=most sociable) using two 4-button devices (Cambridge Research Systems, Ltd., UK). During each trial (Figure 1), a face image was displayed on the screen and participants had 4 s to respond to the image. After responding, the rating was displayed for 0.5 s in the middle of the screen followed by a random intertrial interval (ITI) of 0.5-2 s. Then, a cue image indicating the feedback group (peer or expert) was displayed on the screen for 1 s. The group rating was then displayed for 1.5 s, with an upward arrow indicating a higher rating than the participant’s, a downward arrow lower, and a percent symbol for an equal rating. Participants first completed 10 practice trials. The session lasted 35 minutes.

**Figure 1.**
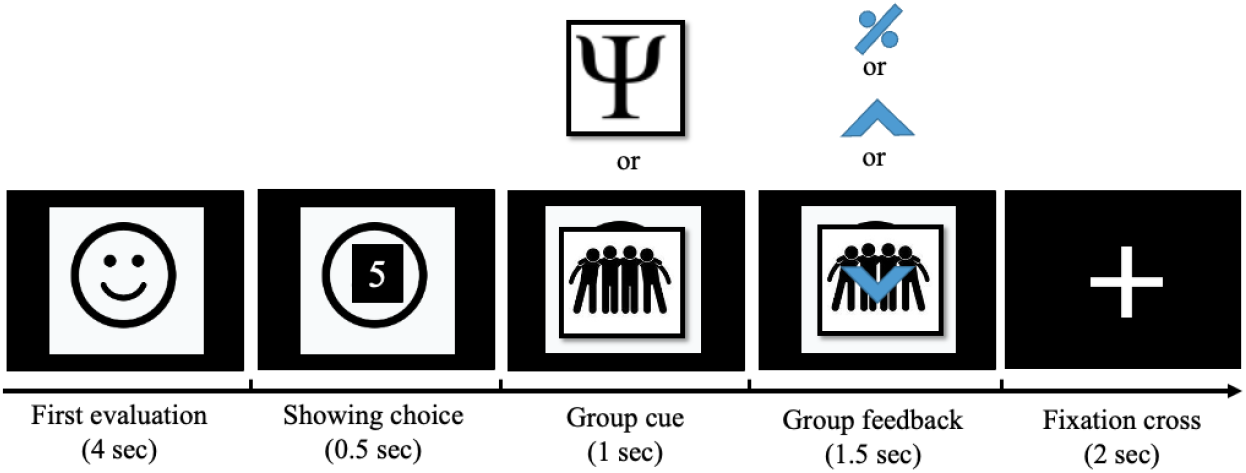
Experimental task design. During the first MEG session, at the beginning of each trial, subjects rated the sociableness of faces on an 8-point scale (actual face was presented instead of the cartoon-face). Then, subjects were presented with a cue image representing either a peer or expert group followed by the presented group rating. Group rating could be the same as the subject (no conflict with group ratings with percent symbol), and different from the subject (conflict with group ratings with upward arrows for positive conflict and downward arrows for negative conflict). Thirty minutes after the first MEG session, subjects rated the same items again without group cues and feedback to identify conformity effects.

After a 30-minute break, we tested if the subject’s initial evaluation had changed after exposure to conflicting group opinions. This second rating session was conducted without prior notice and the faces were rated in a random order without group cues or feedback.

The feedback ratings were scripted using Presentation software (Neurobehavioral Systems, Inc., Albany, CA, USA) to either match the participant’s rating (no-conflict condition) or to be higher or lower (positive and negative conflict condition). The subjects were told they would participate in a study on ‘first impressions’ and evaluations of peer and expert groups have been collected earlier. The peer group would consist of hundreds of high school students including their schools, and the expert group of psychologists from Finland. Therefore, they were unaware of the real aim of the experiment. After the experiment, subjects were interviewed, and the true nature of the experiment was revealed. No participant guessed the study was about social influence.

To investigate whether personality traits mediate the effects of group opinion on conformity behavior, participants were asked to complete several questionnaires. All participants completed the Finnish version of the behavioral activation system/behavioral inhibition system (BAS/BIS) (Carver & White, 1994), Interpersonal reactivity index (IRI) (Davis, 1980), and Peer Pressure Inventory (Short) questionnaires (Santor et al., 2000). A manipulation test questionnaire was administered to determine whether participants observed differences between the peer and expert groups and their feedback.

Seven days after the experiment, all participants were invited to take part in a follow-up online experiment to test the longevity of the conformity effect. 20 out of 23 re-rated the same 240 photographs.

### MEG data acquisition

The MEG data were collected using the TRIUX MEG device (MEGIN Oy, Helsinki, Finland) located in a shielded room at the University of Jyväskylä. The data were collected with a sampling rate of 1000 Hz and a band-pass filter of 0.1–330 Hz. The participants’ head position was monitored continuously using five head position indicator (HPI) coils. The HPI coil positions and three anatomical landmarks (nasion and left and right preauricular points) were digitized using the Polhemus Isotrak 3D tracker system (Polhemus, Colchester, VT, United States). Electro-oculogram (EOG) was recorded using electrodes above the right and below the left eye to capture blinks and saccades and the electrocardiogram (ECG) with electrodes on the left and right clavicle. The ground electrode was attached to the right clavicle bone.

### Data analysis

#### Behavioral data analysis: Conformity

To assess whether individuals shifted their opinions toward peer and expert group’s opinions between the first and second rating sessions, we grouped the trials into three categories: negative, positive, and no-conflict, based on the groups’ opinion compared to the subject’s opinion (Figure 2(a)). A linear model with a complex method was used to assess opinion change, with parameters estimated using MLR estimation in Mplus v8.2 (Muthén & Muthén, 2017). Intraclass correlations (ICCs) were calculated to determine the proportion of the variance in opinion change at within and between-subject levels. Based on calculated ICCs (ICCs change =0.018), the hierarchical nature of the data (n=5492 trials nested within 23 participants) was taken into account by estimating unbiased standard errors using the TYPE = COMPLEX option in Mplus. The null hypotheses about the parameter estimates were first tested using a Wald test. Subsequently, the three categories were converted into dummy variables, allowing us to contrast the negative and positive conflict with the no-conflict (d1 and d2 respectively, Figure 2(c)). The initial (first session) rating was included as a covariate to control for the regression to mean (RtM) effect caused by repeated measurements (Yu & Chen, 2015). We examined the main effect of the feedback group on rating change as well as the interaction between the feedback group and feedback. Upon finding no significance of these factors on rating change, we removed them from the final model.

**Figure 2.**
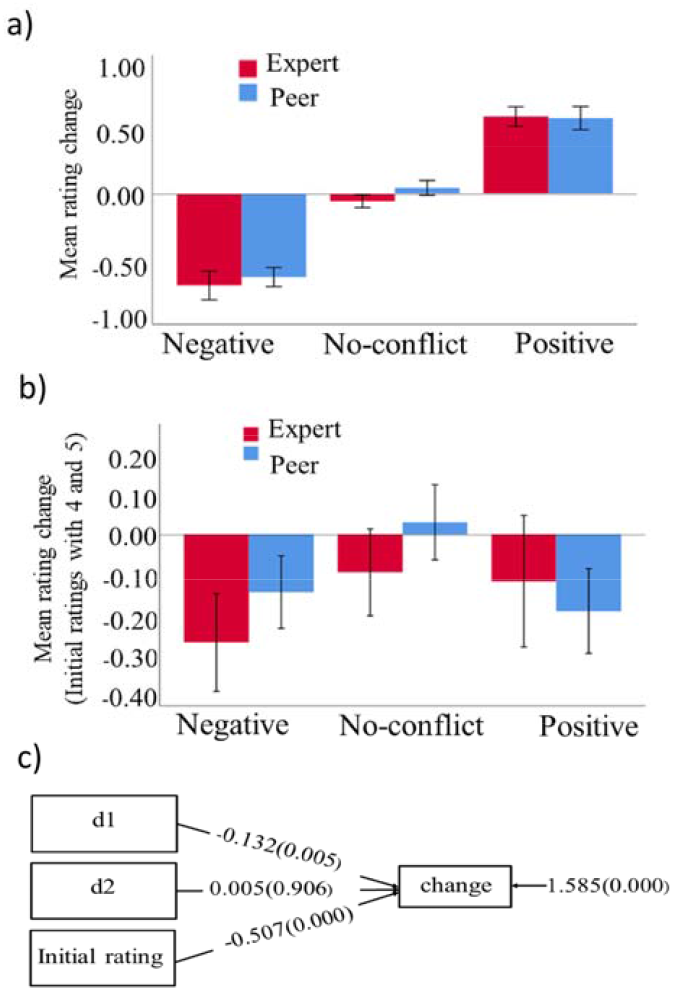
Behavioral results. (A) Mean rating change (second rating minus first rating) for peer and expert groups with all trials across all 23 subjects (total number of trials: 5492). (B) Mean rating change when the effect of the regression to mean (RtM) is removed by using only initial ratings 4 and 5 (2277 trials). Bars indicate the standard error of the mean across subjects. (C) The diagram for statistical analysis shows the behavioral effects of positive and negative conflict and the initial rating. The values indicate mean differences between negative conflict with no-conflict (d1), positive conflict with no-conflict (d2), and the effect of initial rating on change. The value on the right indicates the residual variance of change, and p-values are shown inside brackets.

#### Behavioral data analysis: experimental manipulation test

A manipulation test was used to evaluate the efficacy of the manipulation of normative and informational influence from the two groups. 5 pairs of questions with ratings 1-7 were designed to assess the normative influence and 4 pairs for the informational influence. The analysis was conducted using SPSS (version 26). To determine the internal consistency and reliability of the questions, Cronbach’s Alpha was calculated for each section: peer normative, expert normative, peer informational, and expert informational. Individual questions were averaged into a single value when the alpha value was above the predetermined threshold of 60%. Paired sample t-test was applied to compare the mean differences between the peer and expert groups for normative and informational influence.

#### MEG data analysis: Preprocessing

To reduce interference and compensate for head movements, the tSSS method from the Maxfilter software (MEGIN, Helsinki, Finland) was employed (Taulu & Hari, 2009). Head position was estimated in 200 ms time windows with 10 ms intervals for movement correction, and the data were transformed to the median head position calculated across all sessions. Raw MEG data (filtered at 40 Hz low-pass and 0.5 Hz high-pass) was subjected to independent component analysis (Fast ICA; Hyvärinen & Oja, 2000) to manually identify and remove artifacts related to horizontal saccades, blinks, and cardiac activity. The subsequent MEG data processing steps were performed in MNE-Python, v0.17 (Gramfort et al., 2013). After ICA, the data were downsampled to 333.33 Hz to reduce data size. The continuous data were segmented into epochs, timed to the presentation of the group feedback (Figure 1), to examine the event-related neural responses. The epochs were grouped into positive, negative, and no-conflict trials, depending on whether the groups’ ratings were higher, lower, or equal to the participant’s ratings, respectively. Positive and negative conflicts were combined for further analysis of the conflict effect. An epoch was rejected if any magnetometer channel exceeded 5 pT or any gradiometer channel 500 pT/m. The trigger-to-stimulus delay, measured with a photosensitive resistor, was subtracted from each epoch.

#### MEG data analysis: Event-related field (ERF)

For the ERF analysis, the continuous MEG data were low-pass filtered at 40 Hz and high-pass filtered at 0.5 Hz using a zero-phase FIR filter. The epochs for the analysis were -200-800 ms with respect to the presentation of the group feedback, with 200 ms baseline mean correction. Evoked fields were calculated by averaging the epochs within conflict and no-conflict conditions separately for groups.

#### MEG data analysis: time-frequency response (TFR)

The raw MEG data were analyzed for frequency content by extracting 500 ms pre-stimulus to 1500 ms post-stimulus epochs. The mean of the 500 ms pre-stimulus baseline and the evoked response were subtracted from each epoch. The time-frequency decomposition was performed using Morlet wavelets with center frequencies 4-60 Hz with 2 Hz intervals, and the number of cycles was set to half of the center frequency. Each epoch was convoluted with the complex wavelet and then the absolute value was averaged across the epochs to obtain the induced amplitude. The epochs were downsampled by 2 and an extra 600 ms was trimmed from both ends to prevent edge effects. The responses were converted to z-scores subtracting the mean and dividing by the standard deviation of the baseline to reduce the variability between subjects.

#### Statistical analysis

To determine differences in neural responses between conflict and no-conflict conditions and identify time windows of the differences, a non-parametric permutation test with clustering correction for multiple comparisons was applied (Maris & Oostenveld, 2007). This test was conducted on a time window of 0 to 800 ms for the ERF analysis and 0 to 1500 ms for the TFR analysis. The difference between conflict and no-conflict was calculated as the dependent sample’s t-statistic at each sample (time point for ERF or time-frequency point for TFR). Samples with t-statistics exceeding an uncorrected threshold of α = 0.05 were clustered based on spatial and temporal proximity. The cluster-level test statistic was computed by summing the t-statistics of samples within each cluster and using the largest cluster-level t-statistic as the test statistic (Maris & Oostenveld, 2007). A permutation distribution of the cluster-level statistics was generated by randomly exchanging the condition labels between epochs, with positive and negative cluster-level statistics computed for 5000 permutations. The observed cluster-level statistic was then compared to the surrogate distribution to determine the permutation-based p-value.

## Results

### Behavioral results: conformity

Trial categorization was done based on the direction of the group’s opinion (negative, positive, or no-conflict) (Figure 2(a)). To determine if participants’ opinions were influenced by the group’s opinion, a comparison was made between the 1st and 2nd ratings. Results showed a significant effect of feedback on opinion change through a Wald test, with W(2) = 8.240, p = 0.016, rejecting the null hypothesis that feedback has no effect on opinion change. The difference between negative and no-conflict (d1) feedback was found to be significant (mean difference = −0.132, p < 0.005, Figure 2(c)), while there was no significant difference between positive and no-conflict (d2) feedback (mean difference = 0.005, p < 0.906). The impact of the initial rating on subsequent rating change due to the RtM was included as a covariate, with a significant effect found (mean = −0.507, p < 0.000). The main effect of feedback groups and interaction between feedback groups and feedback were not significant and were therefore excluded from the model.

We also conducted both simple and multiple mediator analyses to examine the potential mediating role of personality traits, which were assessed using three self-report questionnaires, on the impact of peer and expert feedback on opinion change. The results indicated a lack of significant direct and indirect effects of the questionnaire measures on behavioral conformity.

Follow-up ratings after 7 days, based on data from 20 subjects, revealed no significant main effect of feedback, feedback groups, or any interaction between feedback groups and feedback on rating change.

### Behavioral results: experimental manipulation test

The five items used to measure normative influence showed strong internal consistency in ratings from both groups (peer group alpha = 0.765, expert group alpha = 0.663). There was a highly significant difference in normative impression between the two groups (mean difference = 0.878 [mean: expert = 3.96, peer = 4.8435], t(22) = -3.647, p < 0.001). The four items used to assess informational influence also had good internal consistency in ratings from both groups (peer group alpha = 0.713, expert group alpha = 0.761). However, there was no significant difference in the informational impression between the two groups (mean difference = 0.391, [mean: expert = 4.76, peer = 4.36], t(22) = 1.408, p < 0.173).

### MEG results: evoked responses

The evoked response, locked to the presentation of groups’ ratings, is depicted in a butterfly plot in Figure 3 for the grand average of both peer and expert conflicts (upper panel), with magnetic fields mapped for the highest peaks. A peak is seen between 200 to 300 ms in the frontal areas. Statistical analysis of the magnetometer data revealed a single cluster where the evoked response to conflicting feedback differed between groups (lower panel of Figure 3). This FRN-like activity appeared in right frontal sensors at 230-400 ms after feedback, showing a higher amplitude for the peer group than the expert group (p < 0.05). Further exploration of our data showed that the difference between the two groups in conflicting feedback arises from their difference in the positive conflict. Analysis of the gradiometer channels did not reveal any difference between conflicting feedback from peers and experts.

**Figure 3.**
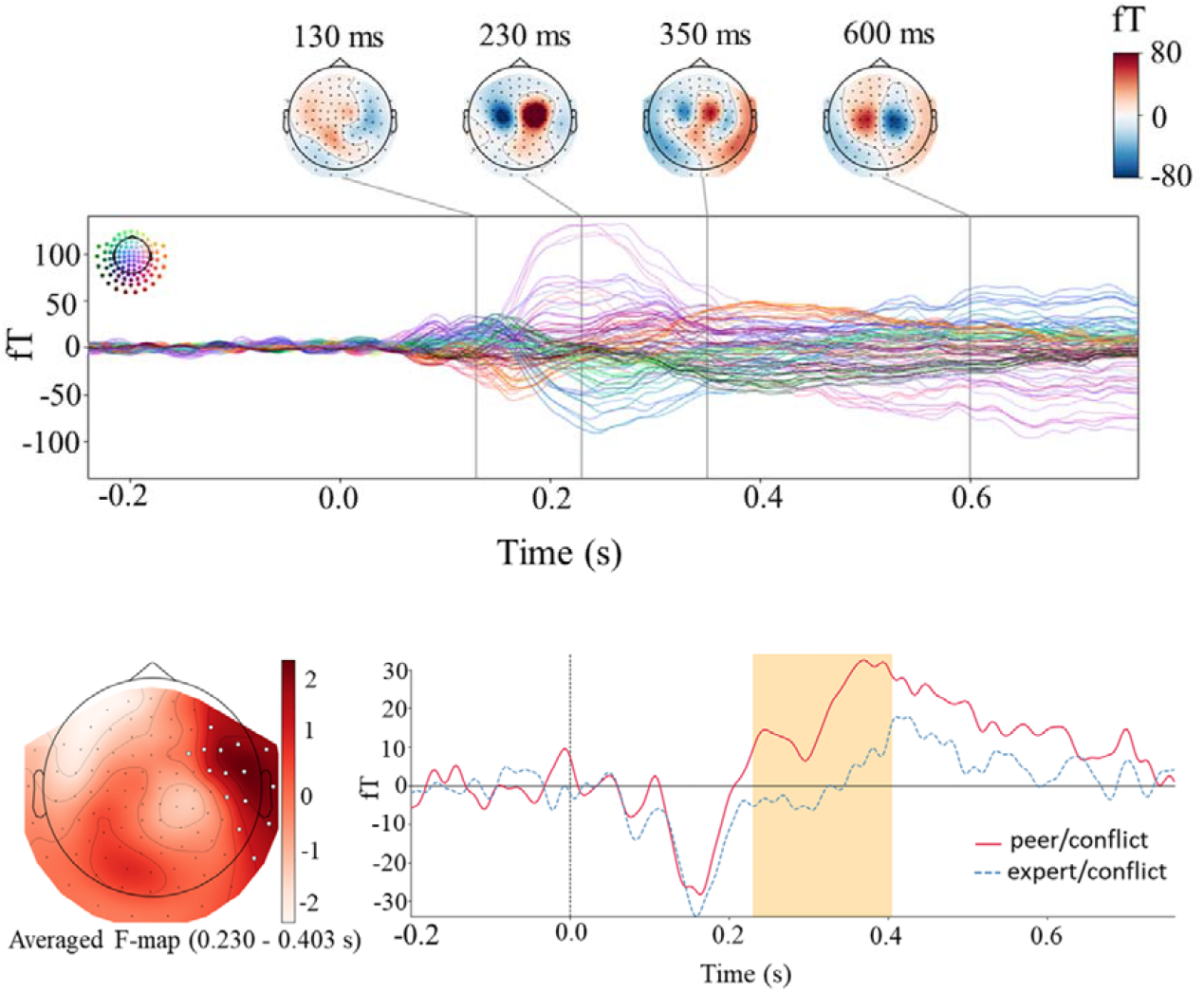
Evoked response analysis. Upper panel. Butterfly plot of magnetometer channels for evoked responses time-locked to the presentation of group feedback for the grand average (peer conflict plus expert conflict). The colored head in the top-left corner shows the individual waveform’s position in the sensor array and the color scales in the top right indicate magnetic field strength. The time points for topographies are selected at activation peaks. Lower panel. Cluster-based permutation test results. Time courses were obtained by averaging over magnetometers comprising the cluster identified by the permutation test. The orange box indicates the time window in which a statistically significant difference was observed.

### MEG results: Induced oscillatory responses

The TFR analysis focused on induced oscillatory activity in gradiometer channels over a time window of -500-1500 ms and a frequency range of 4-60 Hz. The grand average TFR map across gradiometer channels, time-locked to group feedback, is shown in Figure 4 (upper panel). Clear responses in the theta-alpha frequency range (4-15 Hz) between 200 to 400 ms and in the beta frequency range (20-35 Hz) between 400 to 1500 ms were visible. A cluster-based permutation test for condition contrast (conflict minus no-conflict) revealed three significant frequency-specific clusters between 20 Hz to 27 Hz and 100 ms to 1500 ms, with a stronger decrease in induced amplitude for the conflict condition (negative t-values, p < 0.05). The effect was localized to sensors in the right frontal and parietal regions (Figure 4, lower panel). Analysis of the magnetometer channels did not reveal any difference between conflict and no conflict.

**Figure 4.**
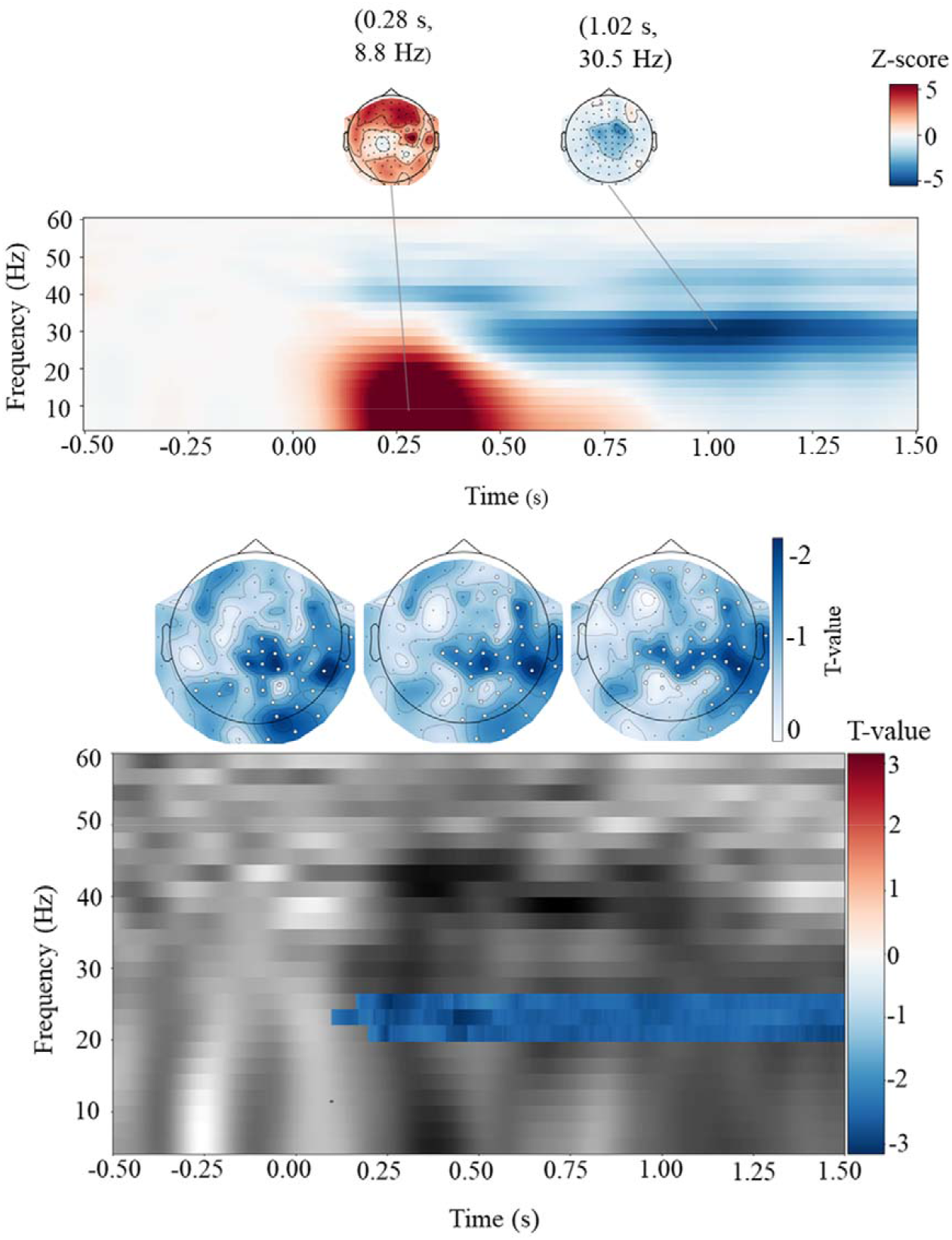
Analysis of induced oscillatory activity. Upper panel. Time-frequency map for grand average (conflict plus no-conflict) with spatial topographies at the peaks. Lower panel. Time-frequency windows and corresponding topographical maps for the three significant clusters resulting from the cluster-based permutation test.

## Discussion

The current study aimed to identify behavioral and neural manifestations of peer and expert group influence on adolescents’ opinion change. Behaviorally, we showed conformity to feedback that was lower than the subject’s rating, but not to more positive feedback. There was no difference between expert and peer feedback on the participant’s opinion change. This finding aligns with prior research indicating that conflicts in a negative direction have a more substantial impact than those in a positive direction (Klucharev et al., 2009; Shestakova et al., 2013; Zubarev et al., 2017; Irani et al., 2022). However, at the neural level, the perceived conflict with the peer opinion evoked greater activity at 230-400 ms time window in the right frontal channels, compared to conflict with the experts. Finally, conflict decreased the power of oscillatory activity in the beta band (20-27 Hz) compared to no-conflict.

Contrary to conventional belief and our hypothesis, peer and expert feedback were not different in influencing opinion change in adolescents in a subjective evaluation task. This is in line with previous studies where feedback from peers and parents induced similar conformity effects in adolescents (Welborn et al., 2016; Do et al., 2020), and even slightly more toward parents (Welborn et al., 2016). Along with the same lines, according to Lourenco et al. (2015), adolescents were found to be influenced by the advice of an older adult, but not by a peer, when they received advice on probabilistic learning choices.

While former studies have suggested that adolescents are particularly susceptible to peer norms (Brown & Larson, 2009; Berns et al., 2010; Blakemore & Mills, 2014; Simons-Morton et al., 2014), our data along with recent findings show that the social contexts under which peer influence has been studied are important. Previous studies on the effect of peer influence during adolescence have typically compared a situation where peers are present to a situation where the individual is alone (Sullivan et al., 2022). Adolescents may conform selectively and even place greater importance on adults’ opinions over peers when comparing the influence of peers to that of other social sources in certain decision-making situations (Wolf et al., 2015).

Our experimental manipulation test indicated that subjects successfully dissociated feedback groups. Moreover, normative influence was observed differently between the peer group and the expert group. In particular, peers were found to have a significant normative influence on participants. Interestingly, both groups inserted equal informational influence on participants. This aligns with Lourenco and colleagues’ (2015) findings that adolescents typically perceive their peers as having expertise in social domains, while experts are viewed as more reliable in non-social contexts such as high-level cognitive tasks (e.g., rational reasoning).

Our follow-up experiment for investigating the longevity of conformity to group opinion revealed no persistent effect of group opinion on preference change. This finding is consistent with Huang et al. (2014), who reported that conformity to group opinion in facial attractiveness ratings lasted only for three days, and frequent daily exposure to faces reset the judgments of attractiveness to the original norm.

At the neural level, conflict with peers evoked more robust activation than conflict with experts in the 230-400 ms time window. The time window during which a difference between peer and expert feedback is observed corresponds to the feedback-related negativity (FRN). The FRN is regarded as a neural indicator of reinforcement learning, specifically negative prediction error, which means learning from errors and conflicts to decrease the likelihood of the behavior being repeated in the future (Shestakova et al., 2013; Zubarev et al., 2017; Irani et al., 2022). FRN reveals temporal aspects of neural activation likely corresponding with the fMRI-registered activation of the medial frontal cortex (MFC) and deactivation of the ventral striatum while viewing conflicting responses from social groups (Klucharev et al., 2009). In our study, the stronger evoked response to peer feedback compared to expert feedback in the FRN time window may reflect that peer feedback elicits a larger prediction error signal. This could be due to the fact that individuals may have stronger expectations about the opinions of their peers and might be more surprised when their opinions differ from peers. On the other hand, expert feedback may be perceived as more objective and less influenced by social factors and norms, leading to a weaker prediction error signal and a smaller FRN. Interestingly, the difference in evoked response between the two groups does not manifest itself in behavior change, as there was no bigger opinion change toward peers. In former probabilistic learning tasks, FRN amplitude correlated with negative prediction error but did not reflect the behavioral adjustment, whereas increased centroparietal positivity (P3 and LPP) were more reliable predictors of behavioral adjustment (Chase et al., 2011; Bogdan et al., 2022). Moreover, Kim et al (2012) found no significant link between error detection and behavior change in social deviance between individuals and group opinions suggesting that detecting an error does not automatically lead to a change in behavior. Shestakova et al (2013) showed that conflict with group opinion elicited an FRN response, indicative of the detection of a violation of group norms. Furthermore, the subsequent positive component was found to be more strongly associated with decision-making processes and behavioral adjustments. Further analysis of our data revealed that the observed difference in evoked response between peers and experts originated from positive conflict, potentially explaining why the difference between the two groups did not manifest itself in conformity behavior.

Our analysis of oscillatory activity showed a decrease in the induced amplitude of beta-band oscillations consistent with other MEG studies on the neural basis of social influence when group feedback differs from one’s opinion (Zubarev et al., 2017; Irani et al., 2022). Previous research suggests that beta-band oscillations are associated with reward signaling and the maintenance of the current sensorimotor or cognitive state (Engel & Fries, 2010). Some studies in feedback learning contexts suggest that there are two types of post-feedback beta activity, with a burst following gains and learning from the positive outcomes and a desynchronization following losses which drives error-based learning and performance improvement (Luft, 2014). The suppression of the beta band in conflict trials here may signal the need for a change in current status and an adjustment of behavior in future encounters.

Former models of decision-making in adolescents suggested that the prolonged maturation of the prefrontal cortex (PFC) results in poor cognitive control on the one hand, and rapid development of limbic regions such as the ventral striatum and amygdala emphasize the role of reward on the other hand, both aspects contributing to characteristics of decision making during adolescence (Steinberg, 2010). Recent studies challenge these theories suggesting that regardless of whether the cognitive control system develops linearly from early to late adolescence or is fully developed by mid-adolescence, adolescents are able to adaptively utilize various forms of cognitive control. Specifically, in social contexts with multiple social sources adolescents effectively engage cognitive control resources to suppress their own antecedent opinions and align their attitudes with others (Welborn et al., 2016; Do et al., 2020; Telzer et al., 2022). In relation to the current study, the manifestation of this flexible cognitive control may be observed through conformity to feedback, irrespective of the group providing it.

To summarize, our study found similar levels of conformity to peers and experts despite heightened neurophysiological responsiveness to the peer group. The inclusion of expert influence may mitigate sensitivity to peers. These findings highlight the need for approaches incorporating other sources of influence alongside peers, rather than solely focusing on changing peer norms to address adolescent behavior.

Our study had a small sample size, with only female participants to avoid cross-gender effects, and further research with larger samples and both genders is needed. Longitudinal research can explore changes in peer and expert conformity patterns across adolescence as our sample included late adolescence. Moreover, the effectiveness of expert feedback may depend on task characteristics, and thus the subjective evaluative judgment task here might not optimally induce the need for specific expertise. Finally, with this particular design, we did not examine the simultaneous influence of peer and expert feedback. In many cases, these sources provide conflicting messages, so it is important to examine how these conflicting messages would affect adolescents’ behavior and brain responses.

## Acknowledgment

The authors thank Kaisa Hytonen for her valuable input in the experimental design.

